# Concentration dependent dual effect of the endozepine ODN on neuronal spiking activity

**DOI:** 10.1101/2022.04.05.487159

**Authors:** M Hazime, M Gasselin, M Alasoadura, D Lanfray, J Leclerc, B Lefranc, M Basile, C Duparc, D Vaudry, J Leprince, J Chuquet

**Affiliations:** Normandie Univ, UNIROUEN, Neuroendocrine, Endocrine and Germinal Differentiation and Communication, INSERM U1239 NorDic, Rouen, France; Normandie Univ, UNIROUEN, Neurological and Ventilatory Handicap Research Group, UR3830-GRHVN, Rouen, France; Normandie Univ, UNIROUEN, Genomic and Personalized Medicine in Cancer and Neurological disorders laboratory, INSERM U1245, Rouen, France; UNIROUEN, Institute for Research and Innovation in Biomedicine (IRIB), Rouen, France

## Abstract

Endozepines, known as the endogenous ligands of benzodiazepine-binding sites, include the diazepam binding inhibitor (DBI) and its processing products, the triakontatetraneuropeptide (TTN) and the octadecaneuropeptide (ODN). Despite indisputable evidences of the binding of ODN on GABA_A_R-BZ-binding sites, their action on this receptor lacks compelling electrophysiological observations, some studies reporting that ODN acts as a negative allosteric modulator (NAM) of GABA_A_R while others suggest the opposite (positive allosteric modulation, PAM effect). All these studies were carried out *in vitro* with various neuronal cell types. To further elucidate the role of ODN on neuronal excitability, we tested its effect *in vivo* in the cortex of the anesthetized mouse. Spontaneous neuronal spikes were recorded by the mean of an extracellular pipette in the vicinity of which ODN was micro-infused, either at high dose (10^-5^M) or low dose (10^-11^M). ODN at high dose induced a significant increase of neuronal spiking. This effect could be antagonized by the GABA_A_R-BZ-binding sites blocker flumazenil. In sharp contrast, at low concentration, ODN reduced neuronal spiking in a magnitude similar to GABA itself. Interestingly, this decrease of neuronal activity by low dose of ODN was not flumazenil dependent suggesting that this effect is mediated by another receptor. Finally, we show that astrocytes in culture, known to be stimulated by picomolar dose of ODN via a GPCR, increased their export of GABA when stimulated by low dose of ODN. Our results confirm the versatility of ODN in the control of GABA transmission, but suggest that its PAM-like effect is, at least in part, mediated via an astrocytic non-GABA_A_R ODN receptor.

## Introduction

Pioneer research on the effect of benzodiazepines (BZ) on GABA_A_ receptor (GABA_A_R)function has led to the discovery of endogenous BZ receptor ligands designated by the generic term « endozepines ». Endozepines is a family of peptides including the diazepam binding inhibitor (DBI, also known as Acetyl Co-A Binding Protein, ACBP) and its processing products, the octadecaneuropeptide (ODN), a small peptide of 18 amino-acids (DBI33-50), and the triakontetraneuropeptide (TTN, DBI 17-50) (Tonon et al., 2019). Endozepines are produced by astrocytes (Compère et al., 2006; Tonon et al., 1990; Vidnyánszky et al., 1994) in most brain structures including the cerebral cortex, the hippocampus and the cerebellum (Tonon et al., 2020). Despite compelling evidence of the binding of DBI and ODN on GABA_A_R-BZ-binding sites, there is no direct information on the precise subunit composition of endozepine-sensitive GABA_A_Rs (Barmack et al., 2004; Bormann, 1991; Dumitru et al., 2017; Möhler, 2014; Qian et al., 2008; Tonon et al., 2020). The existence of another type of receptor, that likely belongs to the GPCR family, has been repeatedly proposed to explain the flumazenil insensitive biological action of ODN (Gandolfo et al., 1997; Hamdi et al., 2012, 2012; Leprince et al., 2001; ALQUIER 2019).

The function of endozepine peptides on GABA_A_R has been inferred from numerous neurophysiological (*i.e*. electroencephalography) or behavioral experiments in which *i*)these peptides mimic the effect of other GABA_A_R allosteric modulators, and *ii*) these effects can be blocked by the BZ antagonist flumazenil. However, the direct electrophysiological effect of exogenously applied endozepine peptides on GABA mediated-inhibitory current in neurons has been rarely investigated. The few available experiments on the effect of some endozepines (ODN essentially) on neuronal activity have been carried out in cultured cells or in brain slices using various types of neurons (e.g. embryonic spinal motoneurons, neural progenitors). From these *in vitro* preparations, it was reported that endozepines act as a negative allosteric modulator (NAM) of the GABA_A_-R, i.e. reducing GABA hyperpolarizing current (Guidotti *et al*., 1983; Ferrero *et al*., 1986; Alfonso et al., 2012; Dumitru *et al.,* 2017). This effect was nearly abolished in mutant mice carrying a phenylalanine (F) to isoleucine (I) substitution at position 77 in the N-terminal domain of the γ2 subunit, rendering the GABA_A_R insensitive to BZs (γ2-F77I mice) (Dumitru et al., 2017). Of note, all these studies have used fairly high concentrations of ODN, in the micromolar range. Following another approach presented in a series of recent publications, Christian and colleagues reported a much more nuanced view of the role of endozepines in the modulation of GABA neurotransmission (Christian et al., 2013; Courtney & Christian, 2018). Using *DBI* KO (DBI^-/-^) mice the authors observed that miniature IPSC frequency and amplitude in CA1 pyramidal cells are increased compared to wild-type mice, in agreement with the NAM effect of DBI and ODN described above (Courtney & Christian, 2018). However, an opposite observation was made in granule cells of the dentate gyrus where the loss of DBI decreased miniature IPSC amplitude and increases their decay time, suggesting that in this subregion of the hippocampus, DBI or its processing products act as positive allosteric modulators (PAM) of GABA_A_R (Courtney & Christian, 2018). The regional specificity of the action of DBI (NAM *vs*. PAM) was also suggested by earlier studies in the TRN (Christian et al., 2013). Another discrepancy between the expected NAM action of ODN and its effect on synaptic plasticity is that neurons overexpressing DBI shows a surprising decrease in LTP (finland paper and thesis). To further elucidate the role of endozepine on neuronal GABA inhibition, we tested the effect of ODN *in vivo* in the cortex of the anesthetized mouse. We confirm previous *in vitro* studies showing that at micromolar concentration, ODN acts as a NAM of GABA_A_-R, increasing the frequency of neuronal firing in a flumazenil dependent manner. In sharp contrast, at low concentration (10^-11^M), ODN reduced neuronal firing, an effect insensitive to flumazenil. As ODN is known to activate astrocytes at low concentration in a flumazenil insensitive manner we further explore whether ODN is involved in the control of GABA release/uptake. In agreement with *in vivo* observations, we found that ODN at low but not high concentration triggers GABA release by astrocytes.

## Material and methods

### Study approval

Experiments, approved by the Ethics Committee for Animal Research of Normandy, were conducted by authorized investigators and in accordance with the recommendations of the European Communities.

### Animals

For these studies, 8-12 weeks old (20-25g) male C57BL/6J mice were purchased from Janvier Laboratories. Animals were housed with free access to standard laboratory diet and tap water, under controlled temperature (22 ± 1 °C) and lighting (light from 7:00 a.m. to 7:00 p.m.). Newborn (24–48 h) C57BL/6 mice were used to prepare secondary cultures of mouse cortical astrocytes for in vitro experiments.

### *In vivo* electrophysiology

Under isoflurane anesthesia (2-2,5%) two small holes were drilled over the whisker barrel cortex with the dura matter intact and two glass micropipettes were inserted (one for the recording and one for the micro-injection). After the surgical procedure isoflurane was reduced to 1.1 ± 0.1% for a resting period of 45-60 min followed by the recording period.

All in vivo recordings were done in a Faraday chamber and used an amplifier PowerLab 8/35 (AD-Instrument). Raw data were acquired with Labchart software (AD Instrument).

Extracellular unit (EU) recording.

An ACSF-filled glass-micropipette/AgCl/Ag electrode, 3-6 μm diameter opening, was positioned in cortical layer 4 of the whisker barrel cortex. Reference and ground electrodes (AgCl/Ag coated wire) were inserted into the cerebellum. For ODN microinjection, a second glass micropipette (10μm diameter opening) was placed 30-50 μm away from the tip of the recording pipette. The signal was bandpass filtered at 200-2000 Hz and digitized at 20 kHz for EU. Signals were recorded for 10 min before the intracortical microinjection of 0.5μL of ODN at 10^-5^ or 10^-11^ M and compared to a 10 min period beginning 3 min after the end of the microinjection (3 min). Spike detection and sorting was then performed semi-automatically, using Klusta software suite (Rossant et al., 2016), freely available on the Web (http://klusta-team.github.io).

### Peptide synthesis

Mouse/rat ODN (H-Gln-Ala-Thr-Val-Gly-Asp-Val-Asn-Thr-Asp-Arg-Pro-Gly-Leu-Leu-Asp-Leu-Lys-OH) was synthesized as previously described (Leprince et al., 2001). Flumazenil, a selective antagonist of the benzodiazepine-site of the GABA_A_R was purchased from Sigma-Aldrich and dissolved in sterile HEPES buffer supplemented with KCl (2.5 mmol/L) and NaCl (145 mmol/L) pH 7.4 and DMSO (dilution 1:4).

### Astrocytes culture

Secondary cultures of mouse cortical astrocytes were prepared as previously described (Masmoudi et al., 2005) with minor modifications. Briefly, cerebral hemispheres from newborn C57BL/6 mouse were collected in DMEM/F12 (2:1; v/v) culture medium supplemented with 2 mM l-glutamine, 1‰ insulin, 5 mM HEPES, and 1% of the antibiotic–antimycotic solution. The tissues were dissociated mechanically with a syringe equipped with a 1-mm gauge needle, and filtered through a 100-μm sieve (Falcon). Dissociated cells were resuspended in culture medium supplemented with 10% FBS, plated in 75-cm2 flasks (Greiner Bio-one GmbH) and incubated at 37 °C in a 5% CO2/95% O2 atmosphere. When cultures were confluent, astrocytes were isolated by shaking overnight the flasks on an orbital agitator. Adhesive cells were detached by trypsinization and seeded for 5 min to discard contaminating microglial cells. Then, the non-adhering astrocytes were harvested and plated in 24 wells plates with an average density of 80 000 cells/ml. All experiments were performed on 48-to 72-h-old secondary cultures (80% confluence). In these conditions, more than 97% of cells were labelled with antibodies against glial fibrillary acidic protein (Gach et al., 2015).

### [^3^H]GABA uptake assay

The measurement of [^3^H] GABA uptake by astrocytes was based on the Fattorini et al. (2017) protocol. The plates (12 wells) were sequentially incubated in a control medium (0.6% DMSO), a medium containing a selective GAT-1 inhibitor, *i.e*. Tiagabine (50 μM, Sigma Aldrich), or a selective GAT-3 inhibitor, *i.e*., SNAP-5114 (30μM, Sigma Aldrich), after which ODN (0, 10-6, 10-9 and 10-12M) was added for 15min at 37 ° C. In the subsequent step, astrocytes were incubated with a mixture of 30 μM of GABA and [3H] GABA (40 μCi, specific activity = 25 Ci / mmol, PerkinElmer, 15 min, 37 ° C.). The supernatant is then removed, the reaction halted by washing with PBS (200 μL, 0.1M) and the cells lysed in a lysis solution (200 μL, 2.5 mL Tris, 5 mL SDS 10%, and 42.5 mL distilled water) and mixed with a scintillator liquid (Ultima Gold, 3mL). The final solution obtained is passed to the liquid scintillation counter (ref of the radioactive counter) in order to quantify the radioactivity in the cells.

### GABA release assay

GABA concentrations in the extracellular medium of astrocytes was determined using the Mouse GABA ELISA Kit (Abbexa, Ltd.) according to the manufacturer’s instructions. Absorbance was measured at 450nm with a plate reader (Infinite 200, TECAN).

### Statistics

Data are presented as mean ± SEM. All statistics were performed using Prism software (Graphpad). Normal distribution of the datasets was tested by a Kolmogorov-Smirnov test. A paired Student’s *t*-test was used for pairwise means comparisons. A Wilcoxon signed rank test was used for pairwise means comparisons of data that did not follow normal distribution. An adjusted p value was calculated using the false discovery rate (Benjamini-Hochberg) method when multiple comparison tests were performed.

## Results

In order to probe the effect of exogenous ODN on cortical neurons activity *in vivo*, we designed a protocol in which ODN was administrated into the cortex with limited mechanical perturbation susceptible to create a bias. After a stabilization period of 45 min under 1.1 ± 0.1% of isoflurane with continuous monitoring of the LFP activity, a first period of 10 min, defined as “PRE”, was recorded. We found that 10 min was a necessary albeit sufficient duration to get a reliable baseline of spiking rate (Figure 1D). Then, 0.5 μL of aCSF was slowly (2.8 nL/s) pressure ejected through a glass micropipette, the tip of which (3 to 6 μM in diameter) was positioned 30-50 μm away from the recording micropipette (Figure 1A). A second period of 10 min recording (define as “POST”) started at the end of the 3 min infusion period. Using this arrangement and injection parameters, aCSF administration didn’t change the spiking rate between PRE and POST period (4 ± 13% change from PRE-aCSF baseline; n = 9 mice; *P* > 0.05) (Figure 2, A and B). As a positive control of the sensitivity of our assay, we micro-injected 10 mM of GABA. As expected, GABA depressed neuronal activity (−58 ± 10% change from PRE-GABA baseline; n =6 mice; *P* < 0.01) (Figure 2, A and B). The neuropeptide ODN at high concentration (10^-5^ M; ODN^High^) induced an increase of spiking rate, with high inter-experiment variability (110 ± 36% change from PRE-ODN^High^ baseline; n = 19 mice; *P* < 0.05). This effect was abolished by the GABA_A_R-BZ site antagonist flumazenil (FLZ) (7 ± 40% change from PRE-ODN^High^/FLZ baseline; n = 5 mice; *P* > 0.05). At low concentration (10^-11^ M; ODN^Low^), ODN significantly decreased the spiking rate (−51 ± 9% change from PRE-ODN^Low^ baseline; n = 8 mice; *P* > 0.05). This effect was insensitive to flumazenil (64 ± 21% change from PRE-ODN^Low^/FLZ baseline; n = 5 mice; *P* < 0.05; *P* > 0.05 relative to POST-ODN^Low^).

**Figure 1.**
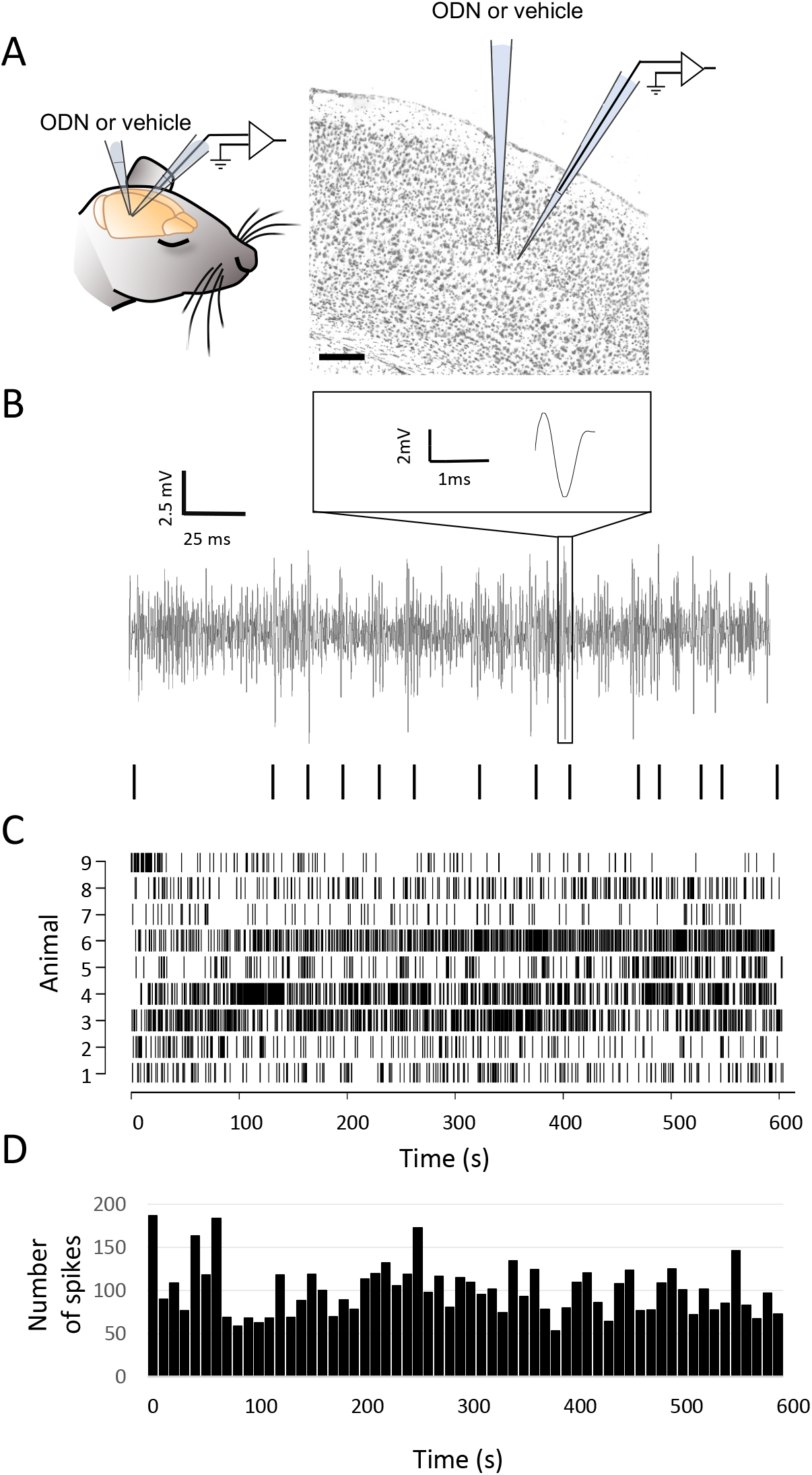

**Figure 2.**
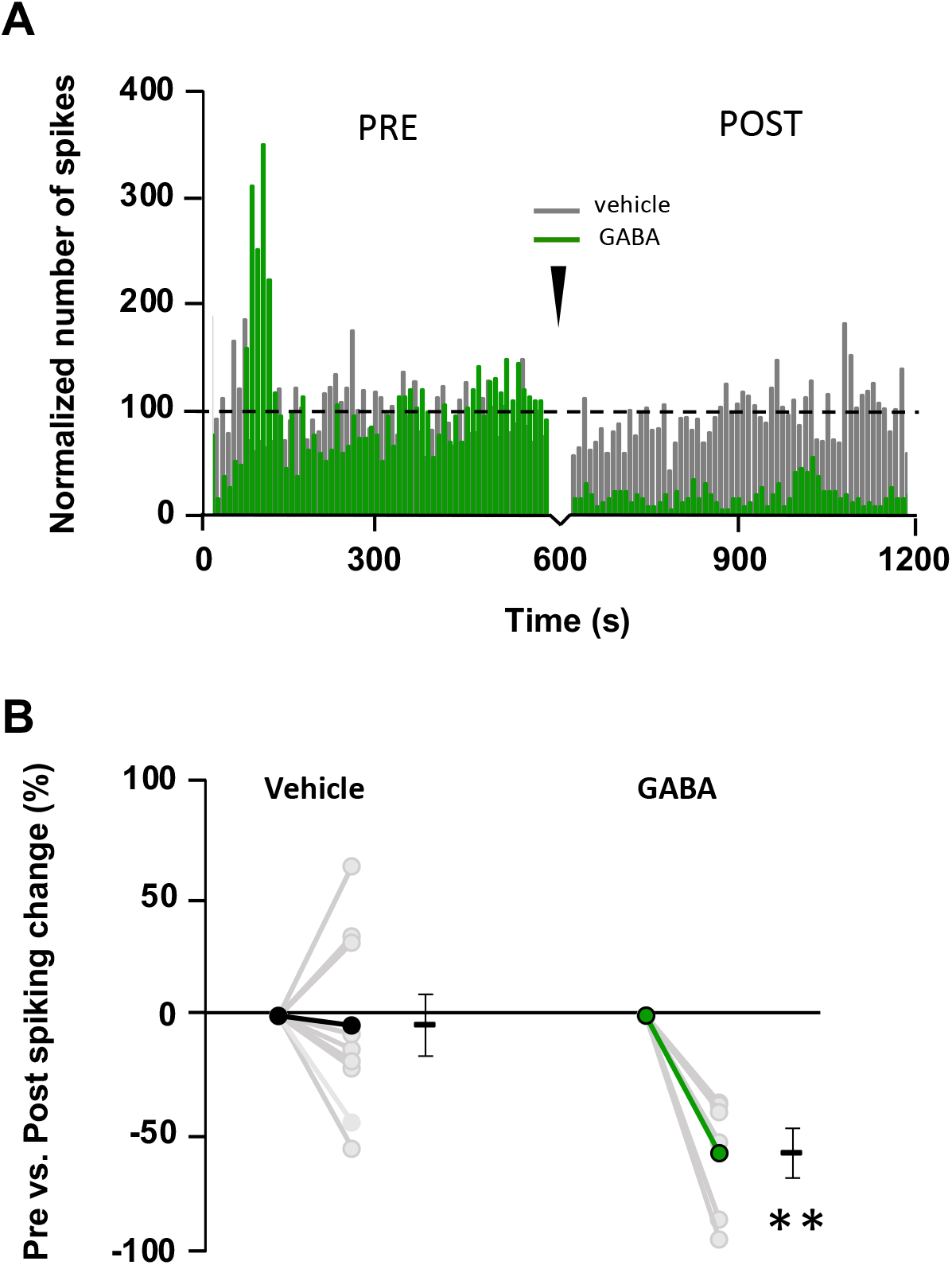

**Figure 3.**
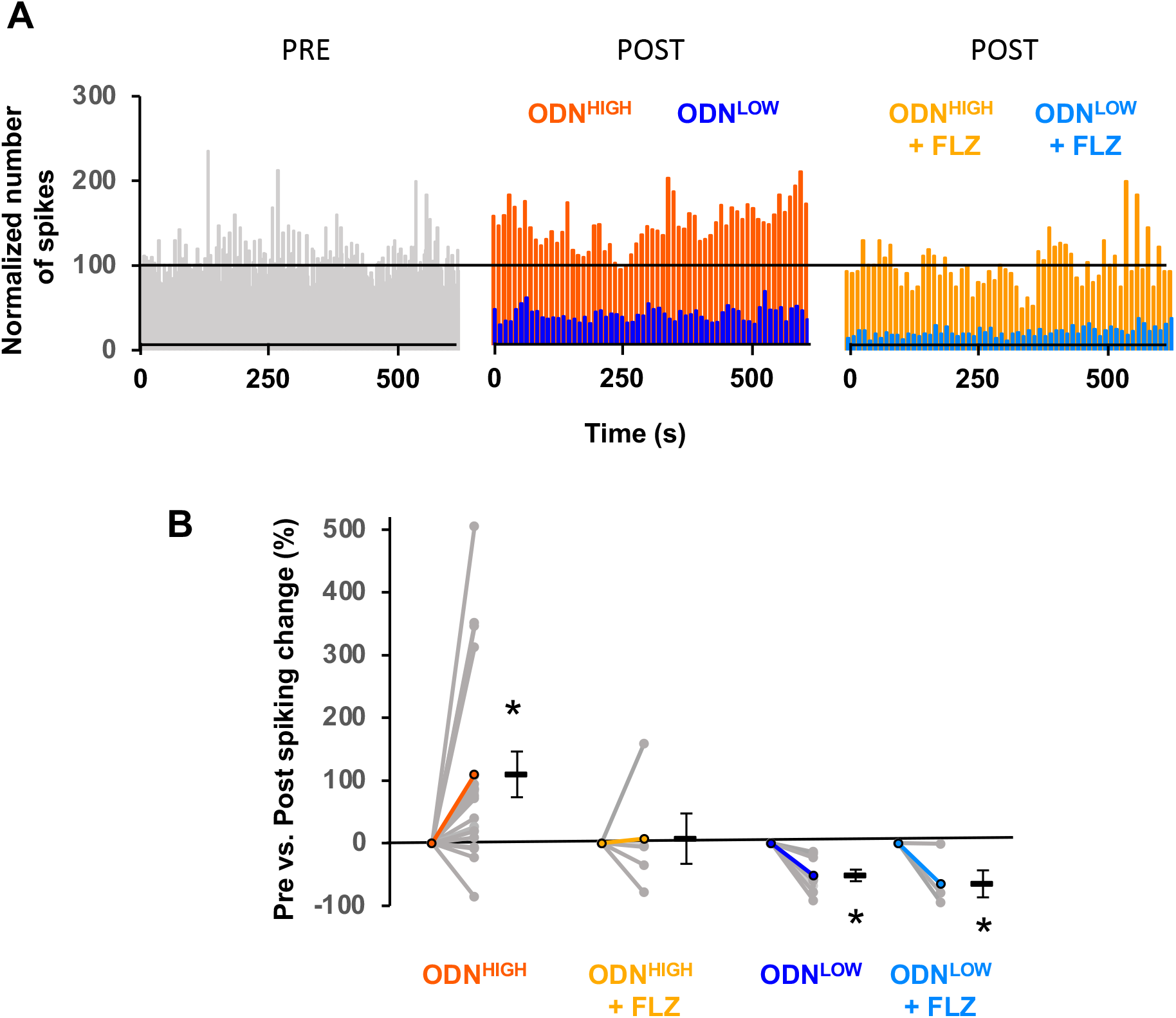

ODN is a gliopeptide exclusively synthetized by astrocytes (Tonon et al., 2019). At low concentration (10^-9^ to 10^-15^ M), ODN is known to stimulate intracellular calcium release of astrocytes in culture (Lamacz et al., 1996), and increase their spontaneous calcium surge frequency both *in vitro* (Gach et al., 2015) and *in vivo* (Lamtahri, 2016). Furthermore, astrocytes are involved in GABA uptake and release in such an extent that it can influence GABAergic transmission (for review, Lia et al., 2019). To test the hypothesis that the ODN-induced neuronal activity change is mediated by astrocytes, we tested the effect of ODN on GABA uptake and release by mouse cortical astrocytes in pure culture.

The influence of *high* (10^-6^ M) and *low* (10^-12^ M) [ODN] was first studied on the release of GABA by astrocytes using an ELISA assay. The infusion of the vehicle medium (100μL) did not elicit any significant change of extracellular [GABA] at any time point (n = 3 cultures with 3 wells/culture for each condition; *P* > 0.05; Figure 4A). Similarly, ODN at high concentration (10^-6^ M) didn’t change the extracellular amount of GABA (n = 3; *P* > 0.05; Figure 4A). In contrast, low [ODN] (10^-12^ M) rapidly induced an elevation of extracellular [GABA], reaching significance at 10 min post-infusion (137 ± 35.4% relative to pre-ODN value; P < 0.05; Figure 4A).

**Figure 4.**
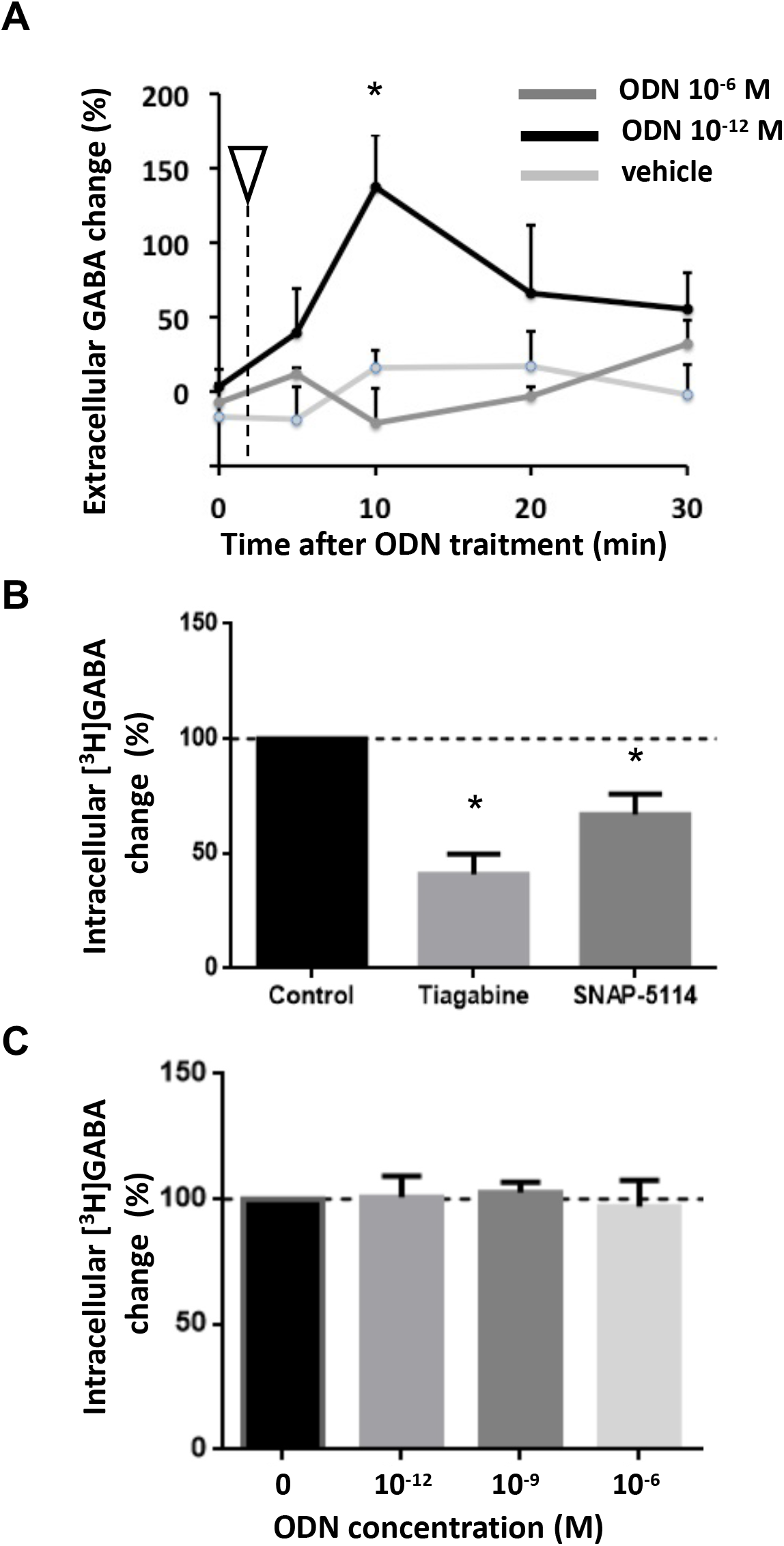

Finally, to address the possibility that ODN modifies astrocytic GABA uptake, we used a radio-assay to quantify the intracellular accumulation of [^3^H]GABA. The uptake of [^3^H]GABA in the presence of the selective GAT-1 inhibitor tiagabine (50 μM) or the selective GAT-3 inhibitor SNAP 5114 (30 μM) was significantly reduced by 59 ± 9 % and 33 ± 9 % respectively (n = 6 and n = 3 respectively; *P* < 0.05; Figure 4B) relative to the total up-take over 30 min in control condition. However, ODN failed to modify the transport of [^3^H]GABA into astrocytes for the 3 concentrations of ODN tested, 10^-6^, 10^-9^ and 10^-12^ M, (n = 4; n = 6; n = 3 respectively; *P* > 0.05; Figure 4C).

## Discussion

The general purpose of the study was to test the effect of the endozepine ODN on neuronal activity, for the first time *in vivo*. Our hypothesis was that the apparent discrepancy reported in the literature about the PAM vs. NAM activity of endozepine for the GABA_A_R may lie in the concentrations at which these peptides operate. In fact, most experiments consistent with a GABA_A_R-NAM effect of ODN have used μM to mM concentration of peptide (Alfonso et al., 2012; Bormann et al., 1985) while many *in vitro* and *in vivo* studies bring evidences that the neuropeptide ODN works at very low concentration, in the fM to pM range (Hamdi et al., 2011, 2012; Kaddour et al., 2019).

The first finding of the present work is that ODN increase or decrease neuronal activity, depending of the concentration used. In agreement with all *in vitro* electrophysiological studies testing the effect of exogenously applied ODN in the micromolar range (no other dosage was actually tried), we found an elevation of neuronal activity, blocked by flumazenil, consistent with a GABA_A_R NAM effect. Surprisingly, we found the exact opposite when ODN was micro-infused at 10pM into the cortex, a reduction of neuronal spiking rate, consistent with the GABA_A_R PAM effect suggested in several recent publications (Christian et al., 2013). It is tempting to believe that, the lowest ODN concentration may be physiologically relevant, while the μM to mM range is never reached in real *in vivo* conditions. However, between brain cells, interstitial volume is extremely narrow (Hrabetova et al., 2018) allowing the concentration of neurotransmitter or other exocytosed molecules to locally reach the mM range. For instance, Overstreet & Westbrook (2003) estimated that GABA reaches a synaptic cleft concentration of 3–5 mM. It is therefore conceivable that ODN, in the close vicinity of its site of release from an astrocytic process, modulate the GABA_A_-R with a NAM effect.

The second important finding of this study is the insensitivity of low concentration of ODN for flumazenil. The decrease of neuronal activity that we observed is unlikely mediated by an interaction of ODN with the GABA_A_R. Indeed, ODN is known to operate at low concentration via a GPCR, the identity of which remains to be elucidated (Bouyakdan et al., 2019; Guillebaud et al., 2017; Tonon et al., 2020). Interestingly, ODN induced calcium elevation in astrocytes is insensitive to flumazenil and has its optimal effect in the pM range. Moreover, at 10^-6^M and above, ODN is no more efficient to induce intracellular calcium transient in astrocytes (Gash et al., 2015). Altogether, these similitudes led us to hypothesize that the observed GABA_A_R-PAM-like effect could actually be the result of an inhibition mediated by astrocytes. There is now a large body of evidence that support the involvement of astrocytes in the modulation of the GABAergic transmission (Lia et al., 2019). In particular, the control of extracellular GABA concentration through the regulation of GABA transport *to* or *from* astrocytes can influence the local neuronal excitability (Woo et al., 2018). To address the relevance of our hypothesis we used an *in vitro* approach with cortical astrocytes in culture. We found that ODN has no effect on the up-take of GABA. Conversely, ODN stimulated the release of GABA, at low but not high concentration. This result supports the view that ODN is not a GABA_A_R PAM, but can nonetheless act to amplify GABA transmission, likely via its GPCR expressed by astrocytes. Further experiments are necessary to validate this hypothesis. Following this idea, ODN at low concentration would enhance astrocyte-mediated tonic inhibition (Farzampour et al., 2015; Tonon et al., 2020).

The function of endozepines remains elusive (Möhler et al., 2014). However, an array of arguments lets us hypothesize that endozepines are instrumental for gliotransmission at the inhibitory synapse. First, DBI is not only specifically produced by astrocytes but is also one of the most transcripted genes in astrocytes (Zhang et al., 2014). Second, potassium depolarization stimulates ODN release from astrocytes (Lamacz et al., 1996). Third, both ODN and GABA provoke an elevation of cytosolic calcium in astrocytes (Gach et al., 2015; Lia et al., 2019). And fourth, extrasynaptic GABA_A_-R are sensitive to BZ and ideally located to bind ODN released by local astrocytic processes. It is therefore tempting to propose a mechanism in which astrocytes sense synaptically released GABA and respond by a release of ODN to modulate positively or negatively tonic inhibition. The present observations make the peptide ODN an appealing candidate as gliomodulator of the inhibitory synapse.

## Aknowledgements

This work was supported by La fondation pour la Recherche sur les AVC.

